# Activity and stress during a critical period regulate olfactory sensory neuron differentiation

**DOI:** 10.1101/2021.01.13.426514

**Authors:** Shadi Jafari, Johan Henriksson, Hua Yan, Mattias Alenius

## Abstract

Here, we reveal that the regulation of *Drosophila* odorant receptor (OR) expression during the pupal stage is permissive and imprecise. We found that olfactory sensory neuron activity directly after hatching both directs and refines OR expression. We demonstrate that, as in mice, *dLsd1* and *Su(var)3-9* balance heterochromatin formation to direct OR expression. Neuronal activity induces dLsd*1* and *Su(var)3-9* expression, linking neuronal activity to OR expression. OR expression refinement shows a restricted duration, suggesting a gene regulatory critical period brings olfactory sensory neuron differentiation to an end. Consistent with a change in differentiation, stress during the critical period represses *dLsd1* and *Su(var)3-9* expression and makes the early permissive OR expression permanent. This induced permissive gene regulatory state makes OR expression resilient to stress later in life. Hence, during a critical period, OR activity feedback similar to in mouse OR selection, defines adult OR expression in *Drosophila*.

## Introduction

Olfactory sensory neurons (OSNs) in most vertebrates and insects are specified to express a single odorant receptor (OR) from a large repertoire of OR genes in the genome (Couto et al., 2005; Fishilevich and Vosshall, 2005; Mombaerts et al., 1996; Ressler et al., 1994). Two OR gene regulatory models have been described: the vertebrate probabilistic selection model and the invertebrate predetermined instructive model.

The vertebrate OR regulatory model depends on chromatin state changes—from a repressed state to an active state and back again to a general repressed state (Lyons et al., 2013; Magklara et al., 2011). In mice, non-expressed OR genes are embedded in constitutive heterochromatin marked by histone H3 lysine 9 trimethylation (H3K9me3) (Lyons et al., 2014; Magklara et al., 2011). According to a mathematical model of OR regulation, a yet-to-be-identified H3K9me3 demethylase sporadically opens the constitutive heterochromatin at a single OR locus and initiates expression (Lyons et al., 2013; Tan et al., 2013). *Lsd1* erases H3K9me2 methylation, which further opens the chromatin and establishes OR expression (Lyons et al., 2013). The expressed OR then induces several feedback loops that down-regulate *Lsd1* and induce heterochromatin formation, blocking the additional initiation of OR expression (Dalton et al., 2013; Ferreira et al., 2014; Fleischmann et al., 2013; Lyons et al., 2013). Unknown transcription factors restrict the expression of each mouse OR to a stereotyped region in the olfactory epithelium (Zapiec and Mombaerts, 2020).

*Drosophila* OR expression is generally viewed as a developmentally predetermined and non-plastic process (Barish and Volkan, 2015; Jafari et al., 2012; Ray et al., 2008). There are several reasons for this assumption. OR expression is stereotypically organized (Couto et al., 2005), and *Drosophila* OSNs are specified in a lineage dependent manner (Barish and Volkan, 2015). Notch signaling splits OSNs into two subgroups with defined projection patterns and OR expression (Endo et al., 2007). Defined transcription factor (TF) combinations both drive and restrict OR expression (Jafari et al., 2012; Komiyama et al., 2004; Sim et al., 2012; Tichy et al., 2008).

Nevertheless, the odor environment and odor exposure early in life can modulate *Drosophila* OR expression and odor responses (Iyengar et al., 2010; Koerte et al., 2018; von der Weid et al., 2015). Thermal stress and starvation induce plasticity in *Drosophila* adult OR expression (Jafari and Alenius, 2015). H3K9me2, which marks OR promoters in vertebrates, also marks OR genes in *Drosophila* OSNs (Sim et al., 2012). *G9a*, which produces H3K9me2, restricts OR expression in *Drosophila* (Alkhori et al., 2014). *Su(var)3-9*, which produces H3K9me3 and induces constitutive heterochromatin, suppresses spurious OR expression (Jafari and Alenius, 2015; Sim et al., 2012). The OR *cis* regulatory regions support cooperative TF interactions that defy heterochromatin and limit stress-induced plasticity (Gonzalez et al., 2019; Jafari and Alenius, 2015). Thus, a heterochromatin regulated OR expression plasticity that is in some ways similar that found in vertebrates also seems to exist in *Drosophila*.

Here, we further address the role of heterochromatin in *Drosophila* OR regulation. We first demonstrate that OR gene regulation stringency increases after a restricted time of heightened plasticity and stress-sensitive period of early fly development. We show that, as they do for vertebrate OR selection, *dLsd1* and su(var) 3-9 initiate and maintain OR expression stringency in *Drosophila*. OSN activity regulates *dLsd1* and *su(var)3-9* expression, creating a feedback loop that restricts and balances OR expression. Stress during this period inhibits the feedback loop and produces permanent changes in OR expression.

## Results

### *Drosophila* chemoreceptor expression matures during the first few days of adult life

We and others have observed that OR reporter expression varies between OSNs in day-old flies, rising to the uniform high level observed in adult flies after a few days (Figure 1A (Golovin et al., 2019)). To further investigate OR expression dynamics, we performed RNA-seq analyses comparing antennae from flies one day (newly hatched), four days (around the point of uniform OR expression), and 14 days (mature) post-eclosion. For each time point biological triplicates were analyzed. Strikingly, all 30 of the 34 adult antennal ORs as well as the olfactory co-receptor Orco increased significantly in expression between one and four days post-eclosion (1-4 DPE, Figure 1B, table S1), but stopped increasing after day four. The expression of the ionotropic receptors (IRs) and gustatory receptors (GRs) also increased during the first four days post-eclosion (Figure 1B, table S1). We found that 13 of 22 antennal IRs and eight of ten GRs expressed in OSNs increased one or more fold. As with the ORs, any changes in IR and GR expression after day four were minor without any discernible pattern (Figure 1B, table S1). Thus, chemoreceptor expression in general seems to mature during the first four days post-eclosion.

**Figure 1.**
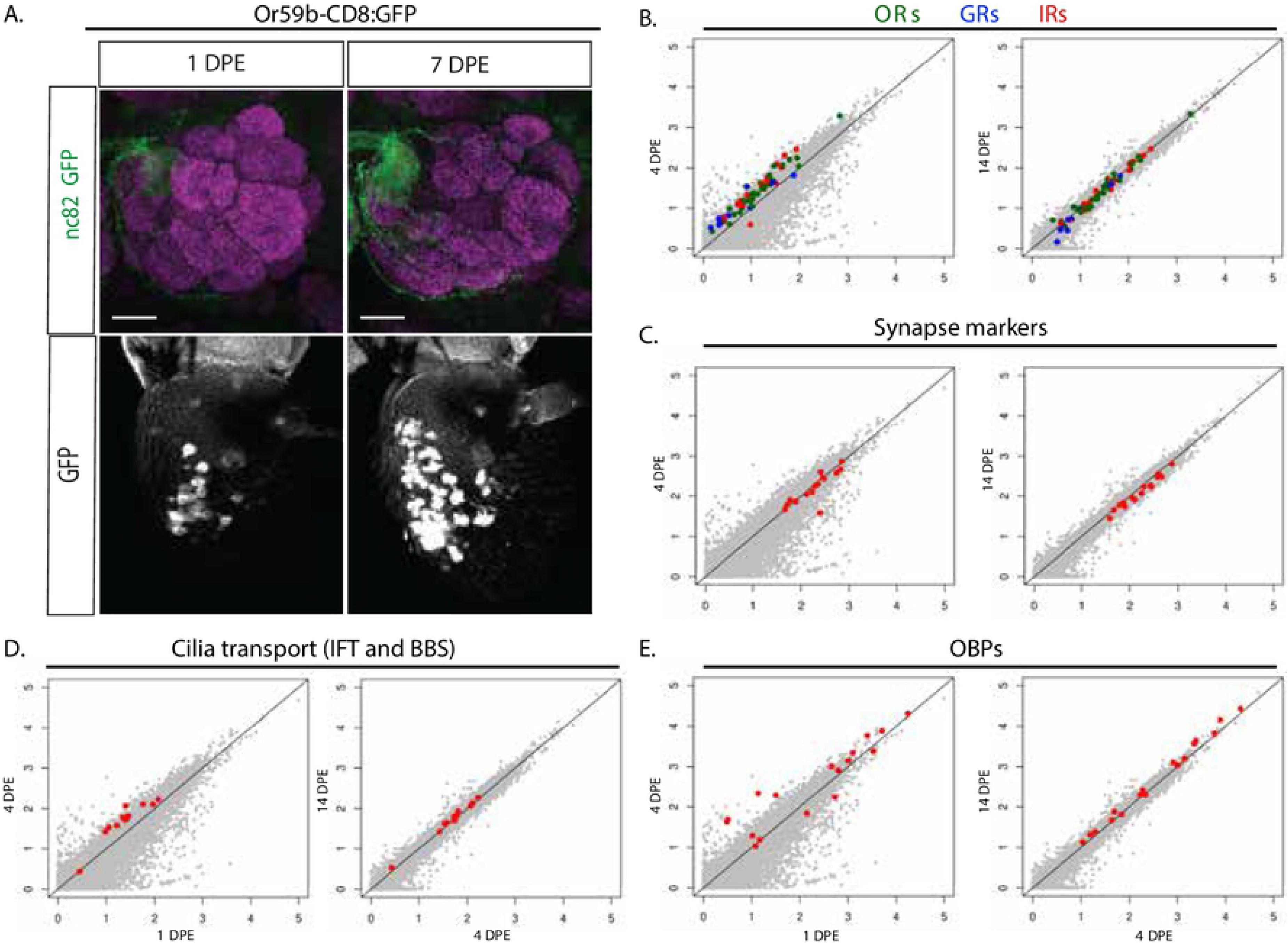
OR expression matures and OSN development continues after eclosion. (A) Whole-mount brain staining and antenna shows the *Or59b* reporter GFP expression (green) in one-and seven-day post eclosion (DPE) flies. Synaptic neuropil regions are labelled with the presynaptic marker nc82 (magenta). Scale bar denotes 3.5 μm. Below each merged image, the GFP expression in the antenna is shown as the white channel. Note the increased expression and uniform level between OSNs in the seven day flies. (B-E) Degree of change in RNA-seq read counts observed between 1 and 4 DPE respective 4 and 14 DPE. Normalized logarithmic counts (log10 size-factor normalized counts) for each gene from the respective sample were scatter-plotted. Genes shown in grey except (B) OR (green), Grs (Blue) and IRs (red) (C) Synapse genes (red), (D) IFT and BBS genes (red), (E) Odorant binding proteins (OBPs) (red). The line is the reference at which gene expression is the same between conditions. Statistics for the figure in Table S1.

During the same period, OSN connectivity is also maturing (Devaud et al., 2003; Golovin et al., 2019; Sachse et al., 2007). Analysis of the synaptic gene network showed a slight decrease but no uniform change in expression of the genes in the synaptic network from day one to day four (Figure 1C, table S1). This indicates that a limited set of genes or a separate post-transcriptional mechanisms are responsible for refining OSN synapses. We next expanded the analysis further to include other OSN gene networks. Sensory neurons are the only ciliated cells in *Drosophila*, and the ciliary transport machinery (e.g., IFTs and BBs) is important for the ciliary localization of the chemoreceptors (Sanchez et al., 2016). In OSNs, olfactory transduction levels are affected by OR levels as more ORs are transported into the cilia (Sanchez et al., 2016). Interestingly, we found increasing expression of the IFT and BB genes during the first four days post-eclosion and no further change after the fourth day (Figure 1D, table S1). OSNs also express high levels of another auxiliary set of olfactory proteins required for specific odor responses the odorant binding proteins (OBPs) (Larter et al., 2016). The expression of most OBP genes either remains steady or increases from day one to day four (Figure 1E, table S1), lending further support to the idea that OSNs continue to develop and sensory transduction continues to change after the pupal stage.

### Stress modulates the maturation of OR expression

We have previously observed that starvation and thermal stress increases OR expression plasticity (Jafari and Alenius, 2015) and that cooperative transcription factor interactions in the *cis* regulatory region stabilize the OR expression. In these studies, we focused on adult (5-7 DPE) flies but the stress induced plasticity suggested that stress could change also the OR expression maturation process. To visualize stress-induced modulation of OR expression at all stages, we used the *Or59b minimal enhancer* (*Or59bME*), an *Or59b* reporter that lacks the cooperative regulation region required to resist stress-induced changes (Jafari and Alenius, 2015). After dissection and whole mount staining of the brain we analyzed the innervation of the antenna lobe. At room temperature (24 degrees), *Or59bME* behaves just like the endogenous *Or59b* gene with its expression restricted to the ab2a OSN class ((Couto et al., 2005; Fishilevich and Vosshall, 2005; Jafari and Alenius, 2015) Figure 2A). Reducing the temperature alters *Or59bME* reporter expression, but the timing of the temperature shift dictates the resulting phenotype (Figure 2). We found that shifts during the first three days post-eclosion led to stereotype ectopic *Or59bME* expression in several OSN classes, as evidenced by the appearance of multiple GFP-positive glomeruli in the antennal lobe (Figure 2A). But all temperature shifts after day three produced loss-of-expression phenotypes (Figure 2B). This sharp transition suggests a drastic change in OSN gene regulation. It also suggests stress may alter terminal OSN differentiation.

**Figure 2.**
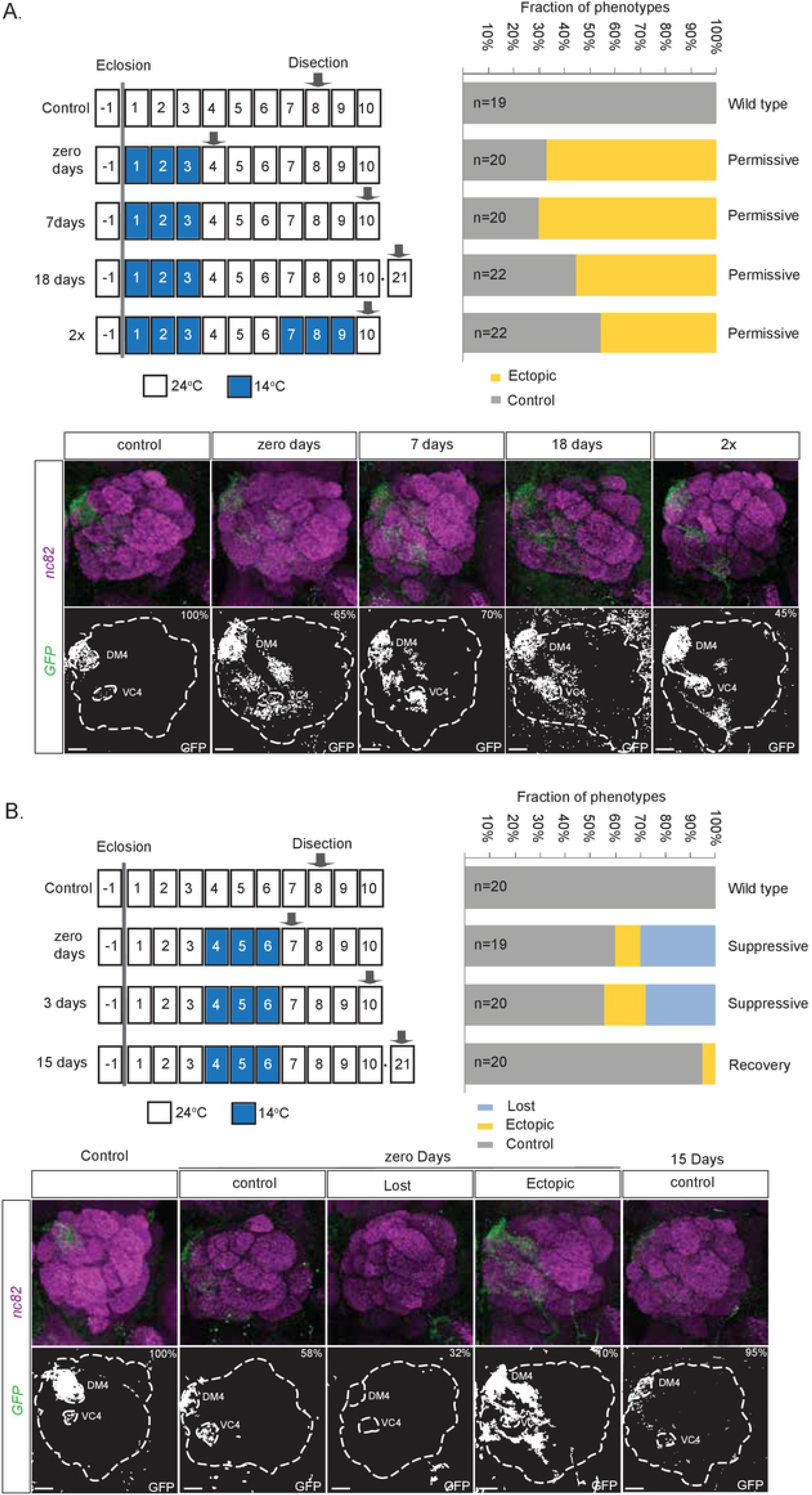
Permanent OR gene regulation changes following environmental stress during the OR expression maturation. The schematic drawing shows the time points of thermal stress treatments and sample preparation. The graphs show the fraction of flies with control, loss or ectopic *Or59bME* expression after three days of thermal stress initiated on (A) day one or (B) day four. The recovery time at ambient temperature denoted in days. The antennal lobes represent the analysed phenotypes. GFP expression (green) is driven by the Or59b minimal reporter. Synaptic neuropil regions are labeled with the presynaptic marker nc82 (magenta). The percentages show the fraction of the phenotype that is presented in that panel. Scale bar denotes 3.5 μm. Note the persistent ectopic expression after 18-day recovery or a second exposure to low temperature (2x). The loss phenotype reverted to single OSN class expression after 14-day recovery at room temperature.

If stress modulates terminal OSN differentiation, the ectopic expression phenotypes we observed with early temperature shifts could be expected to become permanent. Indeed, when we returned *Or59bME* flies that underwent early temperature shifts to room temperature, we found that the stress-induced ectopic *Or59bME* reporter expression pattern persisted throughout a seven-day recovery period (Figure 2A). It even remained similar after a prolonged 18-day recovery period (Figure 2A). If the process of OR expression maturation is the final stage of OSN differentiation, then temperature shifts after maturation is complete should be reversible. Consistent with this hypothesis, we found that shifts back to room temperature for those exposed to thermal stress after day three led to a restoration of the expression pattern in only a single OSN class (Figure 2B). This indicates that the OR expression state was already fixed when the flies were subjected to the temperature shift. To address this further, we subjected flies carrying *Or59b*ME to two cold shifts, one during the critical period and another after a four-day recovery period. As expected, the resulting *Or59bME* ectopic expression pattern for flies subjected to shifts was similar to those subjected to a single early shift (Figure 2A). Together, these results indicate that stress during the maturation phase switches adult OR expression from a stress-sensitive refined expression pattern to a potentially less refined but stress-resilient expression pattern.

### OR activity refines OR expression

In mosquitoes, ectopic OR expression suppresses endogenous OR expression (Maguire et al., 2020). To determine whether OR activity feeds back on OR expression in *Drosophila* as well, we expressed an OR in all OSNs with *Peb-Gal4* and monitored *Or59b-CD8:GFP* reporter expression. With the exception of the male pheromone receptor Or47b, most *Drosophila* ORs have low spontaneous activity (Hallem and Carlson, 2006). We found that about half the ectopic *Or47b* expression flies lost the *Or59b* reporter expression (Figure 3A and B), indicating that high spontaneous OR activity can suppress OR expression. Consistent with the hypothesis that it is OR activity rather than OR expression driving this repression of OR expression, we found that ectopic expression of *Or42b*, an OR with low spontaneous activity, lost *Or59b* reporter expression in 11% of the resulting flies (Figure 3A and B). To determine whether odor responses induce this negative feedback, we exposed flies to ethyl propionate (EP diluted 10^-4^), a strong Or42b ligand. In flies with ectopic *Or42b* expression, EP increased the fraction of flies with loss of *Or59b* reporter expression from 11% to 18% (Figure 3B). EP exposure of control flies (without ectopic *Or42b* expression) did not change *Or59b* reporter expression (Figure 3B). Together the results indicate that the response to environmental odor can shape OR expression.

**Figure 3.**
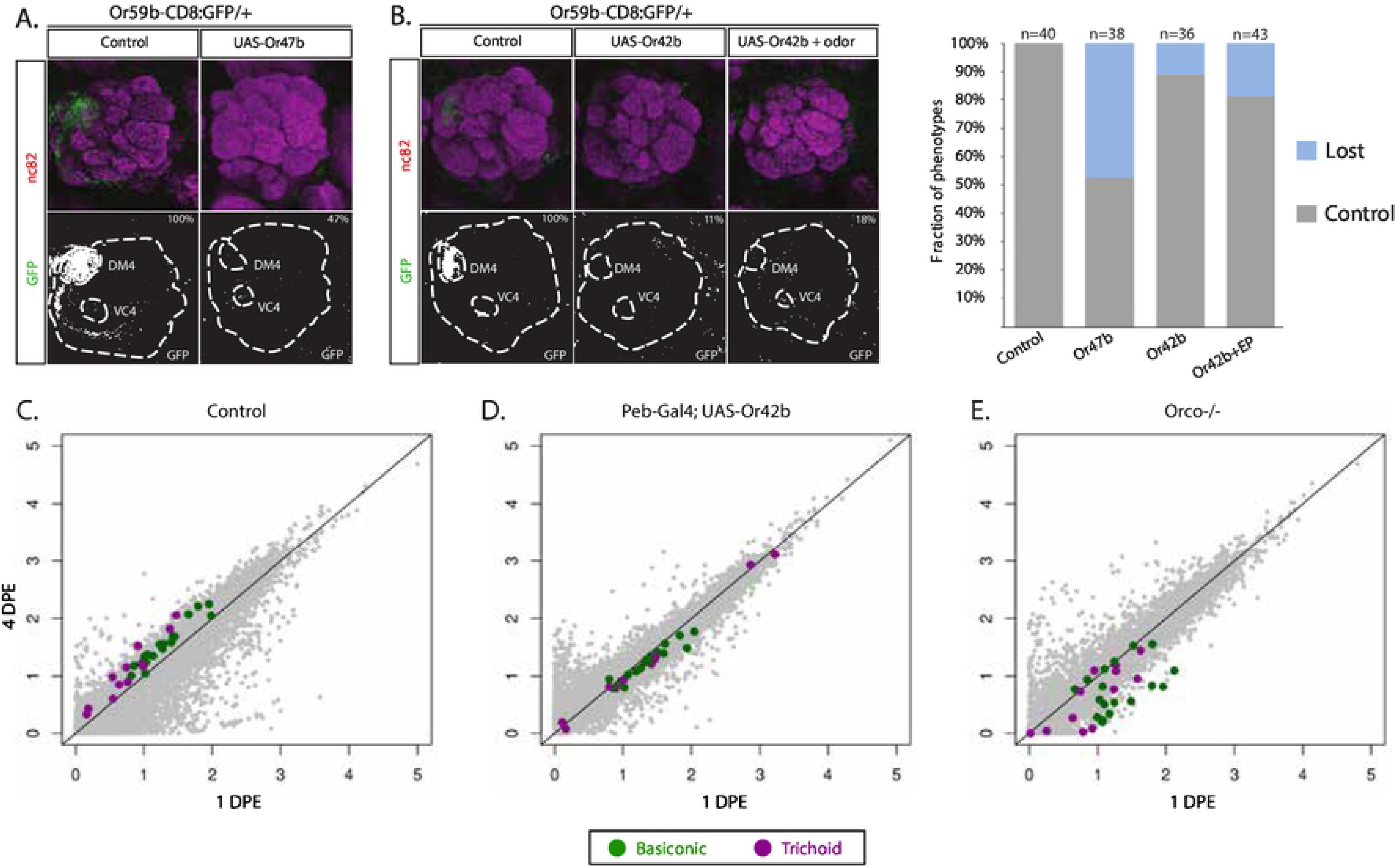
OR activity regulates OR expression. *Or59b* reporter GFP expression (green) and synaptic neuropil regions are labeled with the presynaptic marker nc82 (magenta). Below each merged image, the GFP channel is shown. Antennal lobe and labeled glomeruli are marked. Control flies were crossed to *w^1118^*. % denotes ratio of flies with the depicted phenotype. Scale bar denotes 3.5 μm. (A) ectopic expression of *Or47b* or *Or42b* (B) inhibit *Or59b* reporter expression. The loss of GFP expression is greater when flies with ectopic *Or42b* expression are exposed to the *Or42b* specific odor ligand (EP). (C-E) Degree of change in RNA-seq read counts observed between 1 and 4 DPE. Normalized logarithmic counts (log10 size-factor normalized counts) for each gene from the respective sample were scatter-plotted. Basiconic related ORs (green) and trichoid related ORs (magenta) according to (Couto et al., 2005) other genes in grey. (C) *control*, (D) *peb-gal4; UAS-Or42b*, (E) *Orco*-/-. Note that the increase in OR expression between day 1 and 4, shifts to suppression in OSNs with over activity (D) and lost activity (E). The line is the reference at which gene expression is the same between conditions. Statistics for the figure in Table S2.

Next, we performed an RNA-seq experiment on antennae from flies with ectopic *Or42b* expression. 30 out of 34 antennal ORs expression decreased in the ectopic *Or42b* expression flies between one and four days post-eclosion (Figure 2D, table S2). Comparing OR expression with age matched controls showed that the OSN activity regulated OR expression depended on OSN lineage. In day old (1 DPE) ectopic *Or42b* expression flies most trichoid-related ORs increased in expression (8/12 Table S2 and Figure S1), whereas Basiconic-related OR expression changes were minor. After OR expression maturation (4 DPE), basiconic-related OR expression was down-regulated and the trichoid-related OR expression changes were less penetrant (Table S2 and Figure S1), suggesting that activity establish trichoid-related OR expression during the pupal stage and restricts basiconic-related OR expression post-eclosion.

To further address whether OSN activity is required for OR expression, we performed an RNA-seq experiment on antennae from *Orco* mutant flies, most of the OSNs of which lack OR activity (Larsson et al., 2004). We found a drastic reduction in OR expression between one to four days post-eclosion *Orco* mutant flies (Figure 3E and table S2), indicating that OSN activity is important for OR expression. Consistent with our ectopic expression results, in day old *Orco* mutant flies most trichoid-related ORs were up-regulated while basiconic-related ORs showed no major direction in the changes (Figure S1). At four days post-eclosion most trichoid and all basiconic OR expression was reduced in the *Orco* mutant compared to controls, supporting that activity post eclosion is required in non-stress conditions to establish OR expression. The results also suggest that prior to eclosion spontaneous activity modulates trichoid-related OR expression and post eclosion OSN activity suppress ORs expression in both lineages.

**Figure S1 Difference between OSN lineages in when activity regulate OR expression** Degree of change in sequence counts observed between control and the different genotypes at 1 respective 4 DPE. Normalized logarithmic read counts (log10 size-factor normalized) for each gene from the respective sample were scatter-plotted. Basiconic ORs green and trichoid ORs magenta, other genes in grey. The line is the reference at which gene expression is the same between conditions; increased expression above and suppression below the line. Statistics for the figure in Table S2.

### The balance between *dLsd1* and *Su(var)3-9* refines OR expression

The similarity of the OR activity feedback we observed to the vertebrate OR choice mechanism suggested a conserved OR regulatory mechanism. In mouse OSNs, Lsd1 catalyzes the demethylation of H3K9me2, opening heterochromatin to initiate OR expression (Lyons et al., 2013). To determine whether *dLsd1 (Su(var)3-3)* is also important for *Drosophila* OR expression, we used *Peb-Gal4* to express a UAS-IR (IR, inverted repeats) line specific to *dLsd1* in all OSNs. We found that 16% of the *dLsd1-* depleted flies showed loss of *Or59b* reporter expression (Figure 4A), suggesting *dLsd1* is important in the establishment of OR expression in *Drosophila*. Interestingly, we found that many more *dLsd1*-depleted flies (43%) showed loss of the *Or59b*ME reporter (Figure 4A and B), which lacks cooperative regulation, compared to the *Or59b* reporter. This suggests transcription factor cooperativity and *dLsd1* are both important for OR expression in flies. In mice, one of the few transcription factors known to regulate vertebrate OR expression, Lhx2 (Kolterud et al., 2004), requires cooperativity to maintain OR expression and counteract heterochromatin formation in OSNs (Monahan et al., 2017).

**Figure 4.**
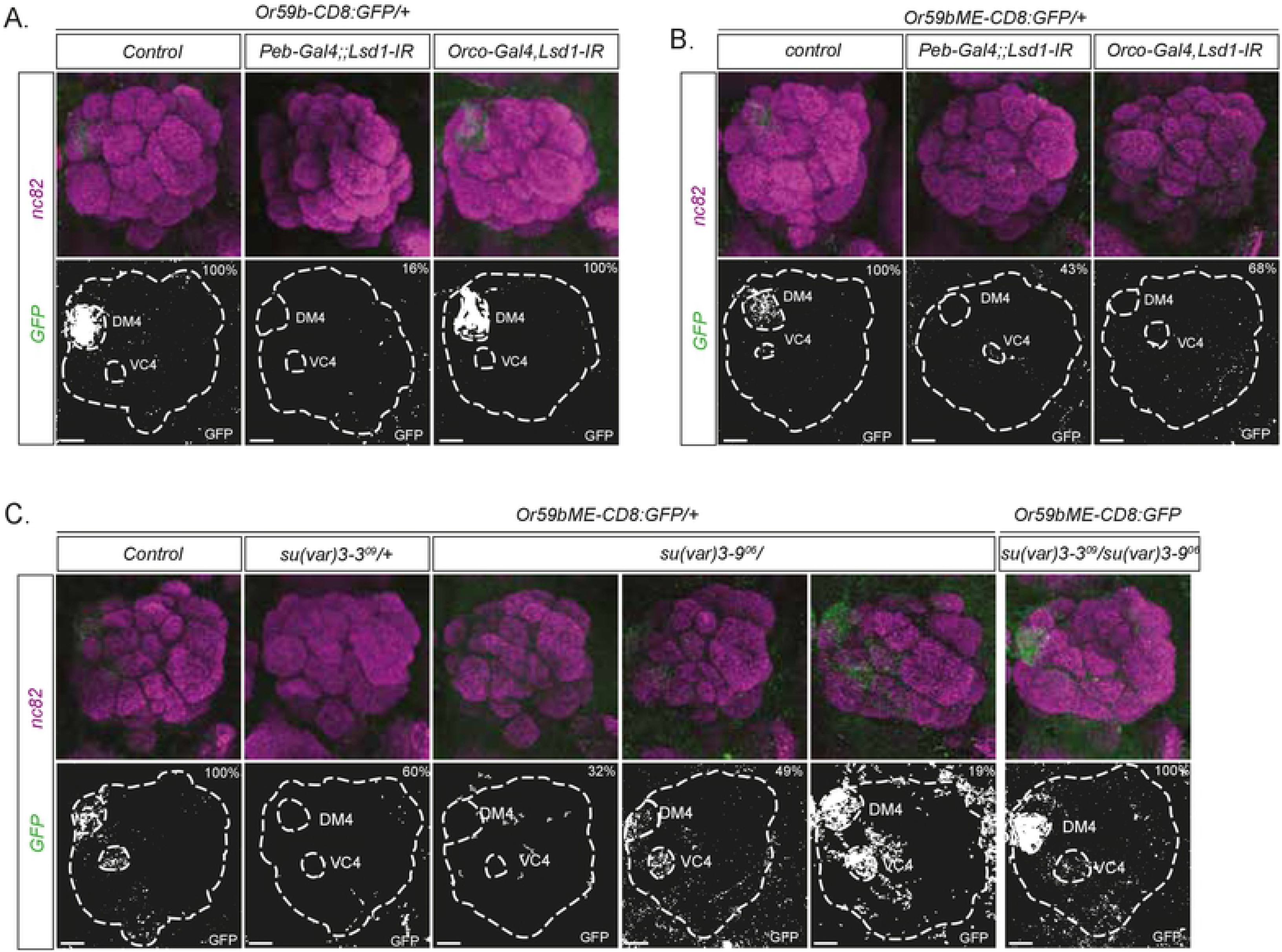
*dLsd1* balance *Su(var)3-9* and establish *Or59b* expression. (A) *Or59b* reporter and (B) *Or59bME* reporter GFP expression in flies with *dLsd1* knockdown before initiation (*peb-Gal4*) or after OR expression (*Orco-Gal4*). GFP expression (green). Synaptic neuropil regions are labeled with the presynaptic marker nc82 (magenta). Scale bar denotes 3.5 μm. (C) *Or59bME* reporter expression in heterozygote *Su(var)3-9^06^*, *dLsd1^09^* and double heterozygote flies. Note that the expression changes of the *Or59bME* reporter in single heterozygote flies was rescued in *Su(var)3-9^06^, dLsd1^09^* heterozygote flies. Control flies were crossed to *w^1118^*. % denotes fraction of flies with the depicted phenotype.

During *Drosophila* development, *dLsd1* erases H3K4me2 and promotes heterochromatin formation (Di Stefano et al., 2007; Rudolph et al., 2007). In both *Drosophila* and in mice, *Su(var)3-9* methylates H3K9me2 to form H3K9me3, a marker of heterochromatin (Nakayama et al., 2001; Rea et al., 2000; Schotta et al., 2002). *Or59bME* reporter expression in heterozygous *Su(var)3-9* mutant flies shows a complex phenotype (Jafari and Alenius, 2015), with 19% of the flies showing ectopic expression, 32% showing loss of expression, and the rest showing single-class expression. *Or59bME* expression is also lost in 60% of heterozygous *Su(var)3-3^09^* (d*Lsd1* mutant) flies (Figure 4C). Combining the *dLsd1* and *Su(var)3-9* heterozygotes resets the balance and rescues reporter expression (Figure 4C). This suggests, not only that the opening and closing of heterochromatin controls OR expression, but also that *dLsd1* promotes open heterochromatin in *Drosophila* OSNs to support OR expression.

To determine whether *dLsd1* initiates or maintains OR expression, we knocked down *dLsd1* using *Orco-Gal4* (Larsson et al., 2004; McLaughlin et al., 2020), which drives expression in most OSNs after OR expression has already begun. In these late knock-down flies, *Or59b* reporter expression was unperturbed (Figure 4A), indicating *dLsd1* is required only during the initiation of OR expression. Interestingly, however, when we repeated the late *dLsd1* knock-down experiment with the *Or59bME* reporter, we found a strong loss of expression phenotype (Figure 4B), showing that *dLsd1* is required continuously to support OR expression.

### *Kdm4b* initiate OR expression

Some mathematical models predict an as-yet-unknown factor that functions at individual OR loci in vertebrates to open constitutive heterochromatin by erasing H3K9me3 (Lyons et al., 2013; Tan et al., 2013). There are two genes encoding H3K9me3 demethylases in the *Drosophila* genome, *Kdm4a* (Kdm4B in vertebrates) and *Kdm4b* (Kdm4A, C, D, E in vertebrates) (Greer and Shi, 2012). We found via knock-down of these two H3k9me3 demethylases in OSNs, that *Kdm4b* but not *Kdm4a* is required for *Or59b* expression (Figure 5A). *Kdm4b* is the major H3K9 demethylase in *Drosophila* (Tsurumi et al., 2013), which is consistent with the hypothesis that the opening of heterochromatin is required for *Or59b* expression. We next asked whether *Kdm4b* is required for continuous *Or59b* expression by knocking down *Kdm4b* after OR initiation. Because OR expression begins in the mid-pupal stage and *Orco* expression begins shortly before eclosion, we decided to use *Orco-Gal4* (Larsson et al., 2004) for this knock-down experiment. We found *Orco-Gal4*-driven *Kdm4b* knock-down had no effect on *Or59b* expression (Figure 5B), indicating that *Kdm4b* is more important for *Or59b* expression initiation rather than maintenance.

**Figure 5.**
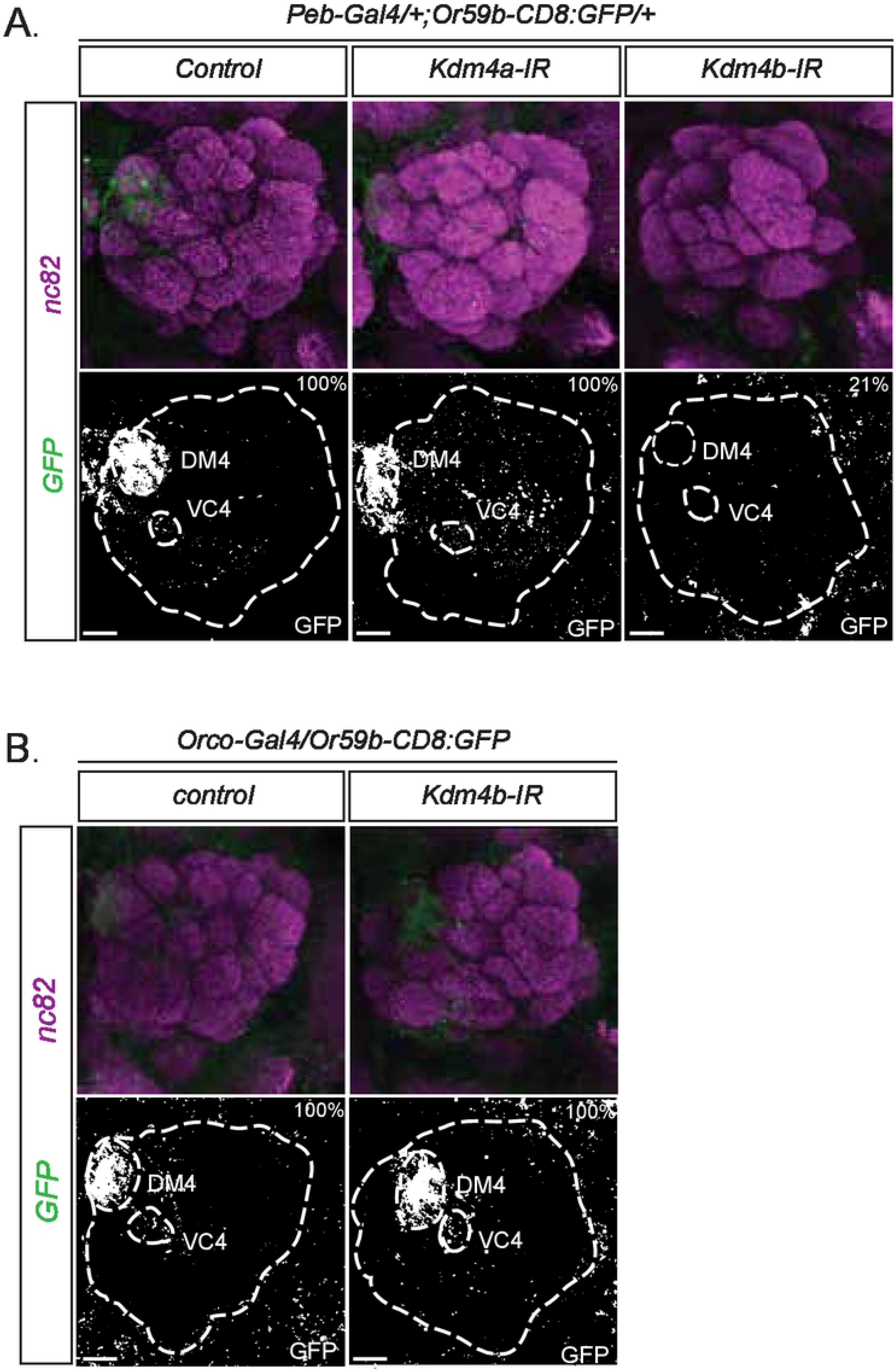
*Kdm4b* initiate *Or59b* expression. Whole-mount brain staining shows *Or59b* reporter expression of GFP (green). Synaptic neuropil regions are labeled with the presynaptic marker nc82 (magenta). Scale bar denotes 3.5 μm. (A) Loss of expression of the *Or59b* reporter is observed in knockdown of *Kdm4b* but not Kdm4a. Control flies were crossed to *w^1118^*. (B) Orco-gal4 knock down of *Kdm4b* after the initiation of OR expression. % denotes ratio of flies with the depicted phenotype. Control flies were crossed to *w^1118^*.

### OR activity regulates *Kdm4b, dLsd1* and *Su(var)3-9* expression

Thus far, our results revealed that the maturation of OR expression comprises a shift from a developmentally permissive state to a more restrictive state in adults. This suggests expression levels of *dLsd1* and *Su(var)3-9* are dynamic. We therefore analysed antennal expression of *Su(var)3-9* and *dLsd1* and found that expression increased from low levels in newly eclosed flies to adult levels three days later (Figure 6A). *Kdm4b* expression, in contrast, decreased over the same period (Figure 6B), suggesting that the maturation of OR expression involves a reduction in the initiation and manifestation of OR expression. A more detailed *dLsd1* and *Su(var)3-9* expression analysis showed that the main increase of these two enzymes occurred during the first hours post-eclosion (Figure 6C), suggesting that OSN activity may induce *dLsd1* and *Su(var)3-9* expression. Consistent with this hypothesis, *Orco* mutant flies that lack OR activity show reduced *dLsd1* and *Su(var)3-9* mRNA levels (Figure 6D). Interestingly, we found Kdm4b mRNA levels also increased in Orco mutants three days post-eclosion compared to controls (Figure 6D). This suggests OR expression initiation likely increases in the absence of OSN activity. Next, we over expressed *Or47b* in a heterozygous *Su(var)3-9* mutant background. We found, consistent with the hypothesis that OSN activity induces heterochromatin formation, that the heterozygote *Su(var)3-9* mutant balanced the ectopic expression of the highly active *Or47b* and rescued the loss of *Or59b* reporter expression (Figure 6E).

**Figure 6.**
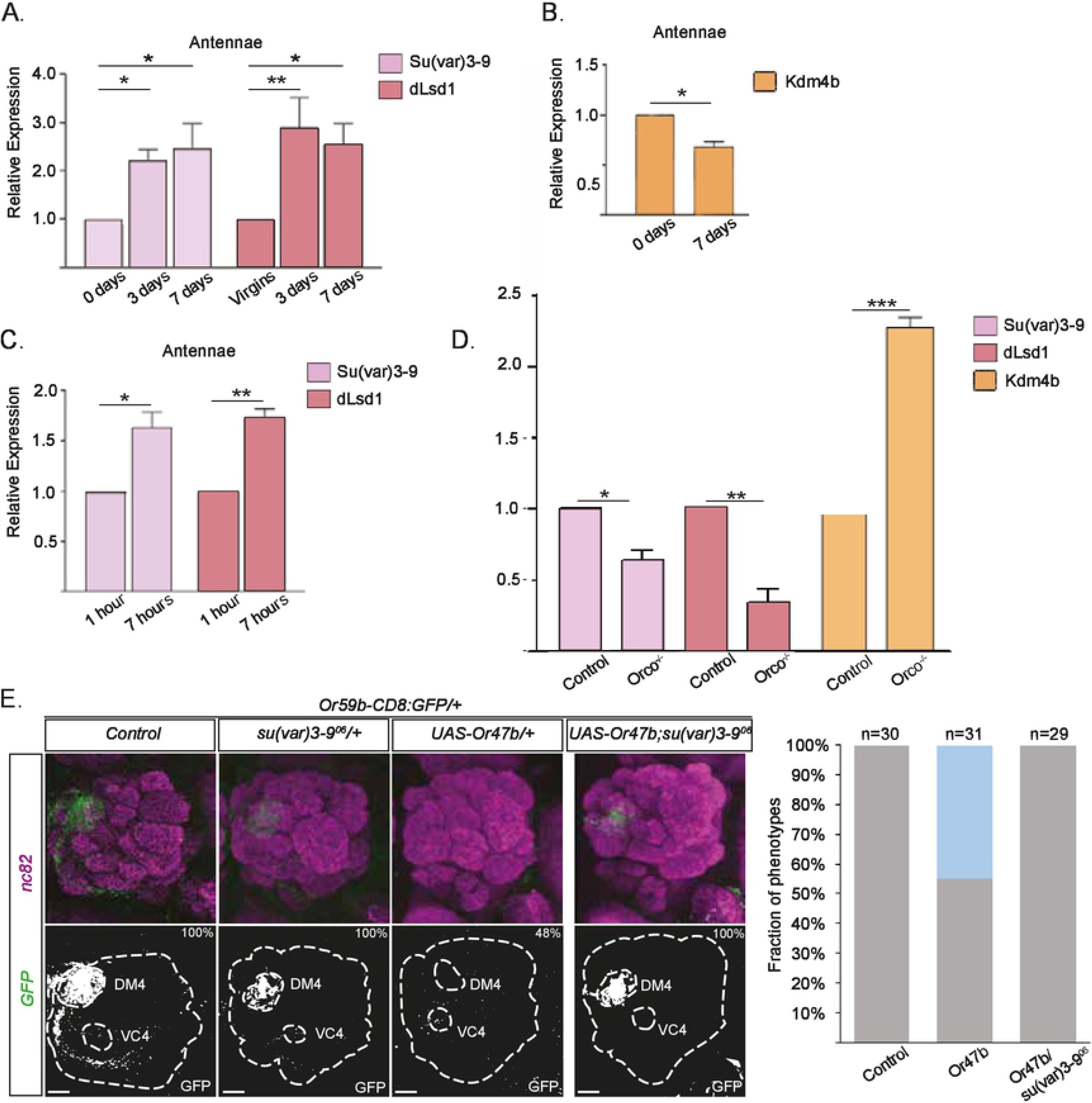
Dynamic expression of chromatin modulators regulates OR expression. (A) The graph shows *Su(var)3-9* and *dLsd1* mRNA levels in antenna at 1, 3 and 7 dpe (* p < 0.05; ** p < 0.01; *** p < 0.001; error bars represent SEM). (B) The graph shows the *Kdm4b* mRNA levels in the antenna at 1 and 7 dpe. Note that *Su(var)3-9*/*dLsd1* show contrasting regulation to *Kdm4b* after eclosion. (C) This graph shows the mRNA levels of *Su(var)3-9* and *dLsd1* in the antenna at 1 hour and 7 hours after eclosion. Note that the expression levels increase to almost twice after the first seven hours post eclosion. (D) The graph compares control (*w^1118^*) and orco mutant mRNA levels of the *Su(var)3-9*, *dLsd1*, and *Kdm4b* in the antenna at 4 DPE. Note that the expression levels are lower for *Su(var)3-9*, *dLsd1* and higher for *Kdm4b* in *Orco* mutant flies. (E) GFP expression (green) driven by the *Or59b* reporter. Note that the loss of *Or59b* reporter expression in Or47b ectopic expression flies is rescued in a *Su(var)3-9*^06^ heterozygote background. The fraction of phenotypes is presented in the graph. Synaptic neuropil is labeled with the presynaptic marker nc82 (magenta). Control flies were crossed to *w^1118^*. Scale bar denotes 3.5 μm.

### Stress regulates *Su(var)3-9* expression differently during and after OR expression maturation

To determine whether *dLsd1* and *Su(var)3-9* expression are sensitive to stress, we analyzed their expression in flies shifted to low temperature at different time points (Figure 7). Flies subjected to a temperature shift at eclosion (1 DPE) showed a two-fold reduction in dLsd1 and Su(var)3-9 expression (Figure 7). The balanced reduction is consistent with a continuous permissiveness. Interestingly, reduction in copy number of both Su(var)3-9 and dLsd1 produced single class expression whereas stress produce ectopic expression, suggesting that additional stress signals enhance the permissive state. After a similar shift in adult flies (7 DPE), *dLsd1* expression fell to the level found at eclosion, whereas *Su(var)3-9* expression showed no significant change (Figure 7). The resulting imbalance between *Su(var)3-9* and *dLsd1* is consistent with the loss of *Or59b* expression we observed when we exposed adult heterozygous *dLsd1* mutant flies to stress (Figure 4C).

**Figure 7.**
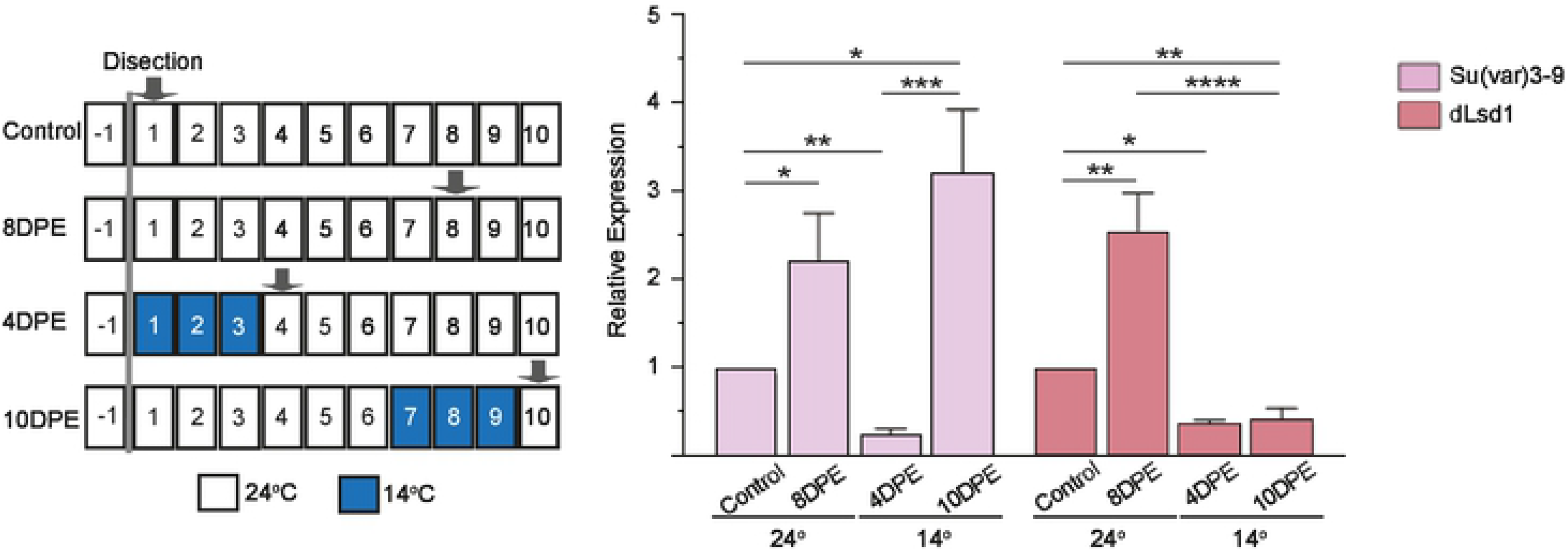
*dLsd1* regulation differs following environmental stress during or after the critical period. Schematic showing the time points of thermal stress treatments and sample preparation. The graph shows the *Su(var)3-9* and *dLsd1* mRNA levels in the antenna after 3 days thermal stress (14°C) ending at 4dpe or 10 DPE, compared to flies maintained at ambient temperature (24°C).

## Discussion

Here, we show that *Drosophila* OR expression matures and that OSNs become terminally differentiated after OR expression initiation. Our results show that OR expression matures in three steps: initiation, establishment, and refinement.

### OR expression initiation: predetermined vs. stochastic

Models of vertebrate OR expression suggest the existence of a heterochromatin switch that initiates expression (Lyons et al., 2013; Tan et al., 2013). We found Kdm4b, an H3K9me3 demethylase, induces OR expression. In a direct instructive model, a predetermined differentiation path produces transcription factors that recruit Kdm4b to open an OR locus. In a more stochastic model, Kdm4b opens chromatin at an OR locus and if the necessary factors are available, the locus is kept open. A recent study revealed that low OR expression precedes the initiation step (McLaughlin et al., 2020), which favors a model in which Kdm4b is recruited to the OR locus or even attracted by low OR expression. We show that OSN activity suppresses *Kdm4b* expression, which suggests that persistent or high OR activity inhibits OR initiation and expression at other OR loci.

### A deeply conserved OR maturation mechanism

The mechanism that establishes OR expression was first identified in mice (Dalton and Lomvardas, 2015). Perhaps the most striking point of conservation is the unique OSN-specific function of *Lsd1*. In most *Drosophila* and vertebrate cells, Lsd1 erases H3K4me1/2 methylation and induces heterochromatin formation (Di Stefano et al., 2007; Rudolph et al., 2007). Our results and several vertebrate OR choice studies (Coleman et al., 2017; Lyons et al., 2013; Vyas et al., 2017) show that Lsd1 opposes *Su(var)3-9* and constitutive heterochromatin formation in OSNs. The enzyme that forms H3K9me2, G9a, also restricts OR expression in both *Drosophila* and mice (Alkhori, Ost, & Alenius, 2014; Lyons et al., 2014), making it clear that H3k9me2 lies at the center of OR gene regulation across phyla. The *cis* regulatory regions have also evolved to balance heterochromatin formation in similar ways between *Drosophila* and mice. Transcription factor cooperativity defies heterochromatin formation and stabilizes *Or59b* expression. In mice, *Lhx2*, one of the few transcription factors known to regulate vertebrate OR expression (Kolterud et al., 2004), also requires cooperativity to block heterochromatin formation and establish OR expression (Monahan et al., 2017).

Also, the feedback loop between OR activity and OR expression was first described in mice (Abdus-Saboor et al., 2014; Dalton and Lomvardas, 2015). The vertebrate feedback mechanisms build on the folding of (Dalton et al., 2013; Lyons et al., 2013) or signaling from an expressed OR (Ferreira et al., 2014; Fleischmann et al., 2013) to inhibit the expression of other ORs. Our result proposes that high OR activity suppress OR expression. We do not explore the art of the activity but that the ORs are ionotropic channels (Benton et al., 2006) suggest a calcium signal to induce the *Su(var)3-9* and *dLsd1*.

After maturation, the OR expression mechanisms differ between *Drosophila* and mice. In *Drosophila*, both OR alleles are expressed in each OSN (Dobritsa et al., 2003). In mice, one OR allele is selected and expressed continuously by what is likely a separate mechanism. Another difference in *Drosophila* is that *dLsd1* activity balances *su(var)3-9* activity after maturation, whereas in mice, Lsd1 is down-regulated after maturation (Dalton and Lomvardas, 2015). We found that it is only *Lsd1* that is supressed after maturation, suggesting the memory mechanism that maintains single OR allelic expression is an inflexible *su(var)3-9* expression that produces a defined heterochromatin level and sets the OR expression baseline. It remains unclear if such a memory mechanism is conserved given the differences in regulation after OR expression maturation.

### A critical period mechanism controls OR expression

The restricted duration of OR expression maturation suggests the period of gene regulation plasticity may be a bona fide critical period (Burggren, 2019; Hensch, 2004). OR regulation does fulfill the criteria. First, a critical period should have a restricted duration, and OR expression maturation ends after a very sharp transition in gene regulation two days post-eclosion. Second, the plasticity of a critical period should be sensitive to neuronal activity, and we showed OSN activity refines OR expression. Third, the phenotype changing in a critical period should be refined through competition, and we showed ectopic OR expression can out-compete endogenous OR expression. Fourth, the plasticity in a critical period should be sensitive to external stress, and we showed that stress can dramatically alter OR expression during and after the relevant period. Fifth, the phenotype developing during a critical period becomes permanent after the period has passed, and we showed that adult OR expression reaches a permanent state, indicating that OSN differentiation ends as the critical period closes. The conserved nature of their mechanisms and the fact that immature vertebrate OSNs also show a low frequency of OR co-expression (Hanchate et al., 2015; Shykind et al., 2004) suggests vertebrate OSN differentiation closes with a critical period as well.

### Differences in activity regulation between OSN classes

We found that the timing of the activity regulation depends on the OSN lineage. According to our results, most of the activity-induced regulation of trichoid-related OR expression takes place in the pupal stage, such that high post-eclosion activity suppresses basiconic-related OR expression much more than trichoid-related OR expression. Interestingly, this difference in regulation relates to olfactory function because basiconic-related ORs respond more to food-related odorants while trichoid-related ORs respond more to pheromones for the sake of social interactions. When a fly emerges from its pupal case, it does so in vicinity of the food it lived on as a larva but not necessarily close to other flies. Consistent with this, pheromone responses increase as social interactions increase post-maturation (Sethi et al., 2019). These response increases come, at least in part, from the sensitization of Or47b OSNs rather than from changes in Or47b expression, suggesting a separate mechanism. Thus, early trichoid-related OR gene regulation supports OR expression even in the absence of stimuli and allows for plasticity even after the OSNs have matured. For basiconic-related OR expression, their late regulation provides more tuning possibilities in a dynamic food odor environment.

### The critical period provides flexibility for OR gene regulation

Predetermined systems of OR gene regulation lacks the flexibility that feedback mechanisms can provide. We and others have shown that high odor responses suppress *Drosophila* OR expression (Koerte et al., 2018; von der Weid et al., 2015). Feedback mechanisms like this tune odor response to environmental odor levels and ensure odor responses fall within physiological limits. Our results further predict that feedback refinement buffers and allows for imperfect gene regulation, reducing the regulatory cost to maintain tight monogenic OR expression. The transcription factors required to express one OR can likely even vary with internal state and stress level. It also predicts that the DNA binding motif locations and *cis* regulatory mechanisms can be plastic between species.

Stress inhibition of the feedback mechanisms also adds to the flexibility of the system. Stress can induce OR paralogue expression, allowing them to contribute to odor responses when the environment changes. In short-lived organisms like *Drosophila*, stress early in life predicts an insecure future. It therefore follows logically that stress early in life would inhibit OR expression maturation, and if the stress lasts beyond the critical period, the changes become permanent. In this way, the permissiveness built into the system makes OSNs and OR gene regulation more robust and resilient to continued or future episodes of stress.

The stress altered OR expression makes also the animal more robust to environmental variability. Our results indicate that the feedback systems and the critical period function as a capacitator, silencing the effect of allelic variability, allowing changes in the OR genes and secure function. This capacitator function hypothesis predicts that in the non-stressed ambient state the activity feedback keeps paralogues and alternate alleles dormant and produce the uniform OR expression observed in adult flies. But when the environment changes, inducing stress, the expression of the dormant OR alleles or paralogues can be activated leading to individualization of the OR expression and the odor responses in the population. Interestingly, the OSNs that express ORs also express IR co-receptors in both *Drosophila* and mosquitoes (Task et al., 2020; Younger et al., 2020), suggesting that ORs and IRs are co-expressed in some OSN classes. Electrophysiology also shown that some OSN classes in *Drosophila* (Ab1b, Ab3a and Ab6a) respond to IR odors (de Bruyne et al., 2001), suggesting that stress can tweak the balance between co-expressed ORs and IRs. Thus, our prediction is that stress accentuate alternative responses and OR allele expression when environmental conditions change and shifts the system from optimal function to maximal detection.

## Materials and Methods

### *Drosophila* stocks

The Or59b promoter fusion and Or59b minimal enhancer constructs were described previously (Jafari and Alenius, 2015). *Pebbled-Gal4* (*Peb-Gal4*) was a kind gift from Liqun Luo (Stanford University, Stanford, CA, USA). The *su(var)3-9^06^* and *Lsd1^09^* mutants were a kind gift from Anita Öst (Linköping University, Linköping, Sweden). The following RNAi lines were obtained from the Transgenic RNAi Project (TRiP; Harvard Medical School, Boston, MA, USA; http://www.flyrnai.org): *Su(var)3-3 (dLsd1)-IR* (36867; 32853, 33726), *Kdm4a-IR* (34629), *Kdm4b-IR* (35676, 57721). The following fly lines were provided by the Bloomington *Drosophila* Stock Center (BDSC; Indiana University, Bloomington, IN, USA; http://flystocks.bio.indiana.edu): *w^1118^* (38690)*, Orco-Gal4* (23909).

### RNAi methodology and environmental experiments

Virgin RNAi females were mated with males carrying *Pebbled-GAL4*, *UAS-Dicer2*, and the cluster transgenes. The crosses were set up and maintained at 24°C. Then, 2–5 d after eclosion, the flies were dissected, stained, and scored for phenotypes.

For the stress experiments, flies were collected as virgins and raised on standard *Drosophila* culture medium at 24°C. On the day for the temperature shift, the flies were transferred to new vials and maintained for three days at 14°C, while control flies were maintained at ambient temperature. Further information can be found in the supplemental experiment statics and details.

### Immunofluorescence

Immunofluorescence was performed as previously described (Jafari et al., 2012). The following primary antibodies were used: rabbit anti-GFP (1:2000, TP-401; Torrey Pines Biolabs) and mouse anti-nc82 (1:100; DSHB). Secondary antibodies were conjugated with Alexa Fluor 488 (1:500; Molecular Probes) and Goat anti-Mouse IgG (H+L) Cross-Adsorbed Secondary Antibody, Rhodamine Red-X (1:250), Thermo Fisher). Confocal microscopy images were collected on an LSM 700 (Zeiss) and analyzed using the LSM Image Browser. The numbers of OSNs co-expressing BP104 and GFP for the different constructs were counted in these images. Adobe Photoshop CS4 (Adobe Systems) was used for image processing.

### qPCR

Antennae were obtained with a sieve after freezing the appropriate flies in liquid nitrogen. Total RNA from the antennae was extracted with TRIzol (Invitrogen) and purified with the RNeasy kit (Qiagen). Quantitative PCR was conducted on an Applied Biosystems 7900HT real-time PCR system (Life Technologies) using the Power SYBR Green PCR master mix (Applied Biosystems, Life Technologies) and primer sets designed using the Primer Express software package v3.0.1 (Integrated DNA Technologies). Actin 5c was used as an internal control. To amplify cDNA products and not genomic DNA, primers were designed to join the end of one exon with the beginning of the next exon. Quantitative PCR for each primer set was performed on both control and experimental samples for 40 cycles. Following amplification, melt curve analysis and ethidium bromide agarose gel electrophoresis were performed to evaluate the PCR products. The relative quantification of the fold change in mRNA expression was calculated using the 2−ΔΔCT threshold cycle method.

### Library preparation

For RNA-seq experiments virgin flies were collected and 50 antennae were handpicked either immediately or after 4 or 14 days on standard *Drosophila* culture medium at 24°C. Total RNA was extracted using TRIzol (Invitrogen, cat. no. 15596-018) according to the manufacturer’s instructions. DNA was degraded using the Invitrogen TURBO DNA-*free*^TM^ Kit. After DNAase treatment, TRIzol RNA extraction was repeated a second time. The concentration and quality of the RNA was determined using a sensitive fluorescence dye-based Qubit RNA HS Assay Kit and the Agilent HS RNA kit and an Agilent 4200 TapeStation System.

Using 1–5 μg total RNA for each sample, we performed two rounds of mRNA isolation using the NEBNext^@^Poly(A) mRNA Magnetic Isolation Module (E7490) according to the manufacturer’s instructions. Libraries were generated using the NEBNext RNA ultra II RNA Library Prep kit. The samples were quality controlled and successfully sequenced on an Illumina NextSeq 500 next-generation sequencing system in mid-output mode via 1X100 bp paired-end sequencing.

### RNA-seq analysis

The RNA read counts were estimated with Kallisto (version 0.45.1). Differentially expressed genes were estimated by DESeq2 (version 1.26.0) after counts had been rounded to the nearest integer count. The linear model was simply one group *vs* the other group, e.g., WT day 1 *vs* day 4, or WT day 1 *vs* Treatment day 1. Plots were made using ggplot2 and R, showing log10 size-factor normalized read counts.

## Data Availability

All the code is available on GitHub (https://github.com/henriksson-lab/mattias-or). The RNA-seq data is available on ArrayExpression with accession #E-MTAB-9805.

## Acknowledgments

We thank Anita Öst and Liqun Luo for flies, Najat Dzaki and Jan Larsson for discussion and comments on the manuscript. Hua Yan acknowledges support from the National Science Foundation I/UCRC, the Center for Arthropod Management Technologies under Grant No. IIP-1821914 and by industry partners. This work was supported by the Swedish Research Council, grant (Vetenskapsrådet VR grant 2016-05208). S.J. is supported by the Swedish Research Council (Vetenskapsrådet VR grant 2016-06726).

## References

Abdus-Saboor, I., Fleischmann, A., and Shykind, B. (2014). Setting Limits: Maintaining order in a large gene family. Transcription 5.

Alkhori, L., Ost, A., and Alenius, M. (2014). The corepressor Atrophin specifies odorant receptor expression in Drosophila. Faseb J 28, 1355–1364.

Barish, S., and Volkan, P.C. (2015). Mechanisms of olfactory receptor neuron specification in Drosophila. Wires Dev Biol 4, 609–621.

Benton, R., Sachse, S., Michnick, S.W., and Vosshall, L.B. (2006). Atypical membrane topology and heteromeric function of Drosophila odorant receptors in vivo. Plos Biol 4, e20.

Burggren, W.W. (2019). Phenotypic Switching Resulting From Developmental Plasticity: Fixed or Reversible? Front Physiol 10, 1634.

Coleman, J.H., Lin, B., and Schwob, J.E. (2017). Dissecting LSD1-Dependent Neuronal Maturation in the Olfactory Epithelium. J Comp Neurol 525, 3391–3413.

Couto, A., Alenius, M., and Dickson, B.J. (2005). Molecular, anatomical, and functional organization of the Drosophila olfactory system. Curr Biol 15, 1535–1547.

Dalton, R.P., and Lomvardas, S. (2015). Chemosensory receptor specificity and regulation. Annu Rev Neurosci 38, 331–349.

Dalton, R.P., Lyons, D.B., and Lomvardas, S. (2013). Co-opting the unfolded protein response to elicit olfactory receptor feedback. Cell 155, 321–332.

de Bruyne, M., Foster, K., and Carlson, J.R. (2001). Odor coding in the Drosophila antenna. Neuron 30, 537–552.

Devaud, J.M., Acebes, A., Ramaswami, M., and Ferrus, A. (2003). Structural and functional changes in the olfactory pathway of adult Drosophila take place at a critical age. J Neurobiol 56, 13–23.

Di Stefano, L., Ji, J.Y., Moon, N.S., Herr, A., and Dyson, N. (2007). Mutation of Drosophila Lsd1 disrupts H3-K4 methylation, resulting in tissue-specific defects during development. Curr Biol 17, 808–812.

Dobritsa, A.A., van der Goes van Naters, W., Warr, C.G., Steinbrecht, R.A., and Carlson, J.R. (2003). Integrating the molecular and cellular basis of odor coding in the Drosophila antenna. Neuron 37, 827–841.

Endo, K., Aoki, T., Yoda, Y., Kimura, K., and Hama, C. (2007). Notch signal organizes the Drosophila olfactory circuitry by diversifying the sensory neuronal lineages. Nat Neurosci 10, 153–160.

Ferreira, T., Wilson, S.R., Choi, Y.G., Risso, D., Dudoit, S., Speed, T.P., and Ngai, J. (2014). Silencing of odorant receptor genes by G protein betagamma signaling ensures the expression of one odorant receptor per olfactory sensory neuron. Neuron 81, 847–859.

Fishilevich, E., and Vosshall, L.B. (2005). Genetic and functional subdivision of the Drosophila antennal lobe. Curr Biol 15, 1548–1553.

Fleischmann, A., Abdus-Saboor, I., Sayed, A., and Shykind, B. (2013). Functional interrogation of an odorant receptor locus reveals multiple axes of transcriptional regulation. Plos Biol 11, e1001568.

Golovin, R.M., Vest, J., Vita, D.J., and Broadie, K. (2019). Activity-Dependent Remodeling of Drosophila Olfactory Sensory Neuron Brain Innervation during an Early-Life Critical Period. J Neurosci 39, 2995–3012.

Gonzalez, A., Jafari, S., Zenere, A., Alenius, M., and Altafini, C. (2019). Thermodynamic model of gene regulation for the Or59b olfactory receptor in Drosophila. PLoS Comput Biol 15, e1006709.

Greer, E.L., and Shi, Y. (2012). Histone methylation: a dynamic mark in health, disease and inheritance. Nat Rev Genet 13, 343–357.

Hallem, E.A., and Carlson, J.R. (2006). Coding of odors by a receptor repertoire. Cell 125, 143–160.

Hanchate, N.K., Kondoh, K., Lu, Z., Kuang, D., Ye, X., Qiu, X., Pachter, L., Trapnell, C., and Buck, L.B. (2015). Single-cell transcriptomics reveals receptor transformations during olfactory neurogenesis. Science 350, 1251–1255.

Hensch, T.K. (2004). Critical period regulation. Annu Rev Neurosci 27, 549–579.

Iyengar, A., Chakraborty, T.S., Goswami, S.P., Wu, C.F., and Siddiqi, O. (2010). Post-eclosion odor experience modifies olfactory receptor neuron coding in Drosophila. P Natl Acad Sci USA 107, 9855–9860.

Jafari, S., and Alenius, M. (2015). Cis-regulatory mechanisms for robust olfactory sensory neuron class-restricted odorant receptor gene expression in Drosophila. Plos Genet 11, e1005051.

Jafari, S., Alkhori, L., Schleiffer, A., Brochtrup, A., Hummel, T., and Alenius, M. (2012). Combinatorial activation and repression by seven transcription factors specify Drosophila odorant receptor expression. Plos Biol 10, e1001280.

Koerte, S., Keesey, I.W., Khallaf, M.A., Llorca, L.C., Grosse-Wilde, E., Hansson, B.S., and Knaden, M. (2018). Evaluation of the DREAM Technique for a High-Throughput Deorphanization of Chemosensory Receptors in Drosophila. Front Mol Neurosci 11.

Kolterud, A., Alenius, M., Carlsson, L., and Bohm, S. (2004). The Lim homeobox gene Lhx2 is required for olfactory sensory neuron identity. Development 131, 5319–5326.

Komiyama, T., Carlson, J.R., and Luo, L. (2004). Olfactory receptor neuron axon targeting: intrinsic transcriptional control and hierarchical interactions. Nat Neurosci 7, 819–825.

Larsson, M.C., Domingos, A.I., Jones, W.D., Chiappe, M.E., Amrein, H., and Vosshall, L.B. (2004). Or83b encodes a broadly expressed odorant receptor essential for Drosophila olfaction. Neuron 43, 703–714.

Larter, N.K., Sun, J.S., and Carlson, J.R. (2016). Organization and function of Drosophila odorant binding proteins. Elife 5.

Lyons, D.B., Allen, W.E., Goh, T., Tsai, L., Barnea, G., and Lomvardas, S. (2013). An epigenetic trap stabilizes singular olfactory receptor expression. Cell 154, 325–336.

Lyons, D.B., Magklara, A., Goh, T., Sampath, S.C., Schaefer, A., Schotta, G., and Lomvardas, S. (2014). Heterochromatin-mediated gene silencing facilitates the diversification of olfactory neurons. Cell Rep 9, 884–892.

Magklara, A., Yen, A., Colquitt, B.M., Clowney, E.J., Allen, W., Markenscoff-Papadimitriou, E., Evans, Z.A., Kheradpour, P., Mountoufaris, G., Carey, C., et al. (2011). An epigenetic signature for monoallelic olfactory receptor expression. Cell 145, 555–570.

Maguire, S.E., Afify, A., Goff, L.A., and Potter, C.J. (2020). A Feedback Mechanism Regulates *Odorant Receptor* Expression in the Malaria Mosquito, *Anopheles gambiae*. bioRxiv, 2020.2007.2023.218586.

McLaughlin, C.N., Brbić, M., Xie, Q., Li, T., Horns, F., Kolluru, S.S., Kebschull, J.M., Vacek, D., Xie, A., Li, J., et al. (2020). Single-cell transcriptomes of developing and adult olfactory receptor neurons in *Drosophila*. bioRxiv, 2020.2010.2008.332130.

Mombaerts, P., Wang, F., Dulac, C., Chao, S.K., Nemes, A., Mendelsohn, M., Edmondson, J., and Axel, R. (1996). Visualizing an olfactory sensory map. Cell 87, 675–686.

Monahan, K., Schieren, I., Cheung, J., Mumbey-Wafula, A., Monuki, E.S., and Lomvardas, S. (2017). Cooperative interactions enable singular olfactory receptor expression in mouse olfactory neurons. Elife 6.

Nakayama, J., Rice, J.C., Strahl, B.D., Allis, C.D., and Grewal, S.I. (2001). Role of histone H3 lysine 9 methylation in epigenetic control of heterochromatin assembly. Science 292, 110–113.

Ray, A., van Naters, W.V., and Carlson, J.R. (2008). A regulatory code for neuron-specific odor receptor expression. Plos Biol 6, 1069–1083.

Rea, S., Eisenhaber, F., O'Carroll, D., Strahl, B.D., Sun, Z.W., Schmid, M., Opravil, S., Mechtler, K., Ponting, C.P., Allis, C.D., et al. (2000). Regulation of chromatin structure by site-specific histone H3 methyltransferases. Nature 406, 593–599.

Ressler, K.J., Sullivan, S.L., and Buck, L.B. (1994). Information coding in the olfactory system: evidence for a stereotyped and highly organized epitope map in the olfactory bulb. Cell 79, 1245–1255.

Rudolph, T., Yonezawa, M., Lein, S., Heidrich, K., Kubicek, S., Schafer, C., Phalke, S., Walther, M., Schmidt, A., Jenuwein, T., et al. (2007). Heterochromatin formation in Drosophila is initiated through active removal of H3K4 methylation by the LSD1 homolog SU(VAR)3-3. Mol Cell 26, 103–115.

Sachse, S., Rueckert, E., Keller, A., Okada, R., Tanaka, N.K., Ito, K., and Vosshall, L.B. (2007). Activity-dependent plasticity in an olfactory circuit. Neuron 56, 838–850.

Sanchez, G.M., Alkhori, L., Hatano, E., Schultz, S.W., Kuzhandaivel, A., Jafari, S., Granseth, B., and Alenius, M. (2016). Hedgehog Signaling Regulates the Ciliary Transport of Odorant Receptors in Drosophila. Cell Rep 14, 464–470.

Schotta, G., Ebert, A., Krauss, V., Fischer, A., Hoffmann, J., Rea, S., Jenuwein, T., Dorn, R., and Reuter, G. (2002). Central role of Drosophila SU(VAR)3-9 in histone H3-K9 methylation and heterochromatic gene silencing. EMBO J 21, 1121–1131.

Sethi, S., Lin, H.H., Shepherd, A.K., Volkan, P.C., Su, C.Y., and Wang, J.W. (2019). Social Context Enhances Hormonal Modulation of Pheromone Detection in Drosophila. Curr Biol 29, 3887–3898 e3884.

Shykind, B.M., Rohani, S.C., O’Donnell, S., Nemes, A., Mendelsohn, M., Sun, Y., Axel, R., and Barnea, G. (2004). Gene switching and the stability of odorant receptor gene choice. Cell 117, 801–815.

Sim, C.K., Perry, S., Tharadra, S.K., Lipsick, J.S., and Ray, A. (2012). Epigenetic regulation of olfactory receptor gene expression by the Myb-MuvB/dREAM complex. Genes Dev 26, 2483–2498.

Tan, L.Z., Zong, C.H., and Xie, X.S. (2013). Rare event of histone demethylation can initiate singular gene expression of olfactory receptors. P Natl Acad Sci USA 110, 21148–21152.

Task, D., Lin, C.-C., Afify, A., Li, H., Vulpe, A., Menuz, K., and Potter, C.J. (2020). Widespread Polymodal Chemosensory Receptor Expression in *Drosophila* Olfactory Neurons. bioRxiv, 2020.2011.2007.355651.

Tichy, A.L., Ray, A., and Carlson, J.R. (2008). A new Drosophila POU gene, pdm3, acts in odor receptor expression and axon targeting of olfactory neurons. J Neurosci 28, 7121–7129.

Tsurumi, A., Dutta, P., Shang, R., Yan, S.J., and Li, W.X. (2013). Drosophila Kdm4 demethylases in histone H3 lysine 9 demethylation and ecdysteroid signaling. Sci Rep 3, 2894.

von der Weid, B., Rossier, D., Lindup, M., Tuberosa, J., Widmer, A., Col, J.D., Kan, C., Carleton, A., and Rodriguez, I. (2015). Large-scale transcriptional profiling of chemosensory neurons identifies receptor-ligand pairs in vivo. Nat Neurosci 18, 1455–1463.

Vyas, R.N., Meredith, D., and Lane, R.P. (2017). Lysine-specific demethylase-1 (LSD1) depletion disrupts monogenic and monoallelic odorant receptor (OR) expression in an olfactory neuronal cell line. Mol Cell Neurosci 82, 1–11.

Younger, M.A., Herre, M., Ehrlich, A.R., Gong, Z., Gilbert, Z.N., Rahiel, S., Matthews, B.J., and Vosshall, L.B. (2020). Non-canonical odor coding ensures unbreakable mosquito attraction to humans. bioRxiv, 2020.2011.2007.368720.

Zapiec, B., and Mombaerts, P. (2020). The Zonal Organization of Odorant Receptor Gene Choice in the Main Olfactory Epithelium of the Mouse. Cell Rep 30, 4220–4234 e4225.

